# Higher-order EEG microstate syntax and surrogate testing

**DOI:** 10.1101/2025.01.07.631820

**Authors:** Frederic von Wegner, Gesine Hermann, Inken Tödt, Inga Karin Todtenhaupt, Helmut Laufs

## Abstract

Higher-order syntax properties of EEG microstate sequences offer insight into the transition dynamics of functional brain networks. We here define higher-order syntax as microstate sequence properties that are not explained by the first-order transition matrix, and we postulate three requirements that surrogate data should fulfill to provide a null hypothesis for higher-order syntax tests. We then compare two general approaches to surrogate data generation that have been used in microstate research, (a) surrogates from a first-order Markov chain model, and, (b) surrogates obtained from sequence shuffling.

There are two different ways of representing microstate sequences, and syntax analyses can be applied to both, continuous microstate sequences, where each time sample is assigned the nearest microstate cluster, or to jump sequences which record only non-identical transitions by removing adjacent duplicates. We show that jump sequences have at least first-order syntax properties, whereas continuous sequences allow for zero-order and first-order surrogates. Markov chain generated surrogates fulfill the three requirements, i.e. they preserve the microstate distribution and transition matrix, and have no higher-order properties. Jump sequence shuffling, on the other hand, yields first-order surrogates whose first-order parameters are markedly different from the original sequence. Using a large open-access resting-state EEG dataset we show that jump sequence shuffling almost certainly produces microstate word probabilities that are significantly different from first-order expected word frequencies, erroneously indicating higher-order syntax properties. Markov chain surrogates reproduce the expected word probabilities of first-order sequences and correctly reject higher-order syntax properties in these cases.

We conclude that jump sequence shuffling does not produce adequate surrogates for higher-order syntax investigations. The proposed Markov chain generative method for surrogate data synthesis is computationally efficient and allows the generation of surrogate sequences of arbitrary length, whereas shuffling can lead to sequences that are shorter than the original sequence and have variable length. Sample code in Python and MATLAB is provided.

## 1 Introduction

Electroencephalography (EEG) microstate analysis is a widely used method for quantitative EEG research (Michel and Koenig, 2018) that compresses and represents EEG datasets by replacing the EEG data vector at a given time with a label taken from a discrete set of labels, or alphabet. These labels represent the cluster centroids obtained from clustering EEG data vectors (topographies) taken at time points where the spatial variance across EEG sensors attains a momentary maximum (Pascual-Marqui et al, 1995; Nagabhushan Kalburgi et al, 2023). In a second step, the entire EEG data set is compressed by competitively fitting the cluster representatives to the multi-channel EEG data vector at each time point, and labeling each time point with the cluster index that gives the best fit (Nagabhushan Kalburgi et al, 2023). Thus, the multivariate EEG time series is reduced to a sequence of discrete labels. Often, the collection of stable EEG topographies is partitioned into four clusters, labelled with the letters A-D, although other choices are becoming more common (Custo et al, 2017). Contiguous blocks of any letter are termed (brain) microstates, and the corresponding topography is called the microstate map. For example, the sub-sequence …ABBBBDA… contains a microstate B with a duration of four samples. Functionally, each cluster can be interpreted as a functional brain network, and the microstate sequence can be interpreted, within limits, as a series of sequentially activating networks (Custo et al, 2017; Michel and Koenig, 2018).

Effectively, the microstate algorithm yields two different but closely related symbolic sequences. The first sequence will be called the continuous sequence in this article, and is obtained from back-fitting the microstate template maps at every time stamp of the discrete EEG signal. Given the average microstate duration ranging from tens to hundreds of milliseconds, and a reasonably high EEG sampling rate, the same microstate label is often repeated across many consecutive time samples in the continuous sequence, thereby producing contiguous blocks. If a given microstate C has a duration of 70 ms, for instance, and the EEG is sampled at 100 Hz (sampling interval 10 ms), there will be a sequence of 7 C’s in the continuous microstate sequence. If we are only interested in the sequence of distinct states and decide to ignore the duration of each state, we obtain the second sequence which is called the (embedded) jump sequence in Markov chain theory, and has been called transitioning sequence (Tait et al, 2020), no-permanence sequence (Artoni et al, 2022, 2023), and compressed sequence (Murphy et al, 2020) in the microstate literature.

The term microstate syntax was introduced in Lehmann et al (2005) where a null hypothesis for bivariate microstate label distributions in microstate jump sequences was introduced. In other words, the probability of microstate blocks of length two (AB, AC, etc.) was described. In terms of Markov processes, the method captures first-order properties, and we will therefore refer to this method as first-order syntax analysis. The same study also identified differences in four-step transitions between individuals with schizophrenia and healthy controls (Lehmann et al, 2005). Many other studies have since used first-order syntax analysis to describe syntax changes under physiological conditions such as sleep (Brodbeck et al, 2012) and development (Tomescu et al, 2018), as well as in numerous pathological conditions such as mood disorders (Al Zoubi et al, 2019), schizophrenia (Tomescu et al, 2015), 22q11 deletion syndrome (Tomescu et al, 2015), sleep deprivation (An et al, 2024), and different forms of dementia (Nishida et al, 2013).

In recent years, there has been a growing interest in the higher-order syntax of microstate sequences where higher-order syntax comprises a range of methods that describe multivariate distributions of microstate labels at different time points (Michel and Koenig, 2018). Moving beyond first-order transitions, Van de Ville et al (2010) investigated fractal properties of continuous microstate sequences, as compared to shuffled sequences, and found significant correlations that often exceed what can be explained by first-order models (von Wegner et al, 2016). We have used higher-order microstate statistics, as compared to Markov chain surrogates, to describe microstate word distributions with information-theoretic and complexity metrics during wake-sleep transitions and anaesthesia (von Wegner et al, 2017, 2018; Wiemers et al, 2023; Hermann et al, 2024). Murphy et al (2020) used sample entropy to quantify the recurrence of microstate words of length 2-10 in early-course psychosis EEG recordings, and used shuffled sequences as surrogates. Artoni et al (2022) quantified the Lempel-Ziv complexity (LZC) of microstate sequences to study propofol-induced loss of consciousness, and later developed a new method for higher-order syntax analysis termed Microsynt that groups microstate words into entropy classes (Artoni et al, 2023). In the latter study, the null distribution was obtained from microstate jump sequences that were shuffled and then corrected for duplicates. The LZC-based study is listed because LZC can be shown to approximate finite-length entropy rate estimates at moderate word lengths. Thus, although the LZC algorithm does not use a fixed word length, it can be seen as a metric of syntactic complexity (von Wegner et al, 2023).

The principal aim of this article is to establish a statistical framework for the systematic testing of higher-order syntax properties with surrogate data, independent of the method that is used to describe higher-order syntax. Instead of evaluating a specific syntax definition, we give a general definition of higher-order syntax within which our considerations are valid. We then compare two approaches for the generation of surrogate data, a) the Markov chain generative method, and, b) the sequence shuffling approach. Both approaches are found in the microstate literature but their differences have not been described and contrasted quantitatively, so far. We characterize both methods first mathematically and then empirically on eyes-closed resting-state EEG data from the public LEMON dataset (Babayan et al, 2019).

This article is structured as follows. We first provide some theoretical background and terminology about Markov processes to make the rest of the article self-contained. This allows us to express the most commonly used null hypothesis for microstate syntax analysis (Lehmann et al, 2005) as a first-order syntax. We generalize the existing definition to give a broad definition of higher-order microstate syntax in terms of microstate word distributions. This definition leads to three requirements that surrogate data should fulfill to be useful for higher-order syntax statistical testing. The two methods for surrogate data generation are then reviewed in detail, and their properties are tested against the three requirements defined before. Python and MATLAB implementations of the Markov chain generative method are included.

## 2 Theoretical background

Microstate sequences will be interpreted as stochastic processes and their syntactic structure will be compared to that of a Markov process of fixed order.

### 2.1 Markov processes

A Markov process of order *m* is a stochastic process (*X*_*t*_)_*t*∈ℤ_ for which an optimal prediction of the future state *X*_*t*+1_ is contained in the last *m* states *X*_*t*_, *X*_*t*−1_, …, *X*_*t*−*m*+1_ of the process (Billingsley, 1961). In the context of microstate sequences, we interpret the discrete variable *t* as time (samples), and use the notation

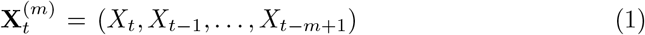

for microstate blocks or words of length *m*. When the focus is on prediction, the term 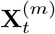 can be called the m-history of *X*_*t*_ (von Wegner et al, 2023). The Markov property of order *m* is expressed as:

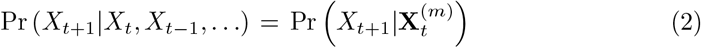

In the following, we will consider microstate sequences over *K* microstate classes and we will use the set of integers ⌊*K*⌋ = {0, 1, …, *K* − 1} as microstate labels. We will sometimes use the microstate labels A-D (Michel and Koenig, 2018) when referring to the literature and implicitly convert to integers 0-3 for array indexing.

#### 2.1.1 Zero-order Markov property

For the Markov property of order zero (*m* = 0), Equation 2 gives

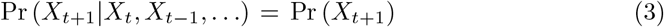

Assuming stationary microstate probabilities during the observation interval, we can ignore the time index *t*, and the best prediction of the next state is defined by the distribution of microstate labels alone. We will write the microstate distribution as **p** = (*p*_0_, …, *p*_*K*−1_). As an example, the short microstate sequence CBACDCDBDC of length *n* = 10 gives **p** = (0.1, 0.2, 0.4, 0.3), as would any other sequence of the same length containing one A, two B’s, four C’s, and three D’s, in any order.

Equation 3 expresses the fact that a zero-order process has no temporal correlations as the next state of the sequence is independent of the sequence’s past values. Testing empirical microstate sequences against a zero-order surrogate therefore only tests whether the process is completely uncorrelated in time. If the zero-order null hypothesis is rejected, there could be first-order (dependence on the last state) or higher-order (dependence on multiple past values) syntactic properties in the tested sequence.

#### 2.1.2 First-order Markov property

A first-order Markov process is defined by setting *m* = 1 in Equation 2:

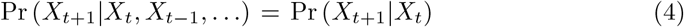

The conditional distribution Pr (*X*_*t*+1_|*X*_*t*_) is called the transition matrix in the EEG microstate literature (von Wegner et al, 2017). For a first-order process, it contains all available information about the transition dynamics. The individual coefficients of the transition matrix T = (*T*_*ij*_)_*i,j*∈ ⌊*K* ⌋_ are the conditional probabilities

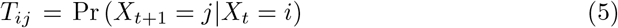

which are obtained as maximum likelihood estimates from the actual microstate counts in empirical microstate sequences (Anderson and Goodman, 1957).

It should be noted that other authors use the term transition matrix for the joint probability distribution Pr (*X*_*t*_, *X*_*t*+1_) (Lehmann et al, 2005). We will use the conditional form because it is commonly used in the Markov process literature, and it is also a practical choice for surrogate data generation, the purpose of this paper (Häggström, 2002). Conditional probabilities for first-order syntax analysis have been used by us (von Wegner et al, 2017, 2018; Hermann et al, 2024) and other groups (Al Zoubi et al, 2019; Antonova et al, 2022; Tomescu et al, 2015) in the past.

#### 2.1.3 Equilibrium distribution

For a full characterization of a first-order Markov process that is not in equilibrium, not only **T**, but also the initial distribution of states **p**_0_ is needed (Häggström, 2002). This initial distribution **p**_0_ = (Pr (*X*_0_ = 0), …, Pr (*X*_0_ = *K* − 1)) contains the probability of each state when the Markov chain is started. For a Markov chain in equilibrium, the transition matrix alone contains all information about the process as the stationary (equilibrium) distribution can be computed from **T**. The equilibrium distribution is defined as a distribution that is not changed by the action of **T** on it, i.e. **pT** = **p**. The distribution **p** can be found as the normalized left eigenvector to the **T**-eigenvalue *λ*_0_ = 1, which always exists since **T** is a stochastic matrix (Perron-Frobenius theorem). In A.2 it is shown that the maximum likelihood estimates for **p** and **T** from any empirical sequence will always fulfill **pT** = **p**, so the non-equilibrium case is not considered further.

A corollary of this observation is that changing the transition matrix **T** is likely to change the corresponding microstate distribution **p** too, resulting in a parameter pair (**p, T**) that describes a different Markov process. This will be investigated in detail further below where we quantify the effects of sequence shuffling on microstate syntax. Another observation is that an uncorrelated (zero-order, syntax-free) process has a transition matrix with a very simple structure. By definition, the zero-order property implies

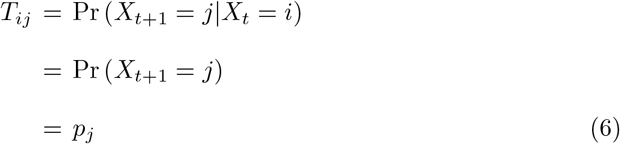

This means that the *K × K* transition matrix of the uncorrelated process has *K* identical rows, and each row is identical to the distribution **p**. The distribution **p** is a valid equilibrium distribution (eigenvector) of this particular form of **T**. We will use this property to describe sequence shuffling, which removes temporal correlations, and to synthesize and analyze surrogates of defined Markov order and arbitrary length.

### 2.2 First-order syntax analysis

The microstate syntax analysis presented by Lehmann et al (2005) is based on the joint probability transition matrix of microstate jump sequences, where consecutive microstate labels must be different. Their null hypothesis postulated that co-occurrences of microstates *i* and *j*, expressed by *P*_*ij*_ = Pr (*X*_*t*_ = *i, X*_*t*+1_ = *j*), are determined by the relative frequency (coverage) of *i* and *j* in the microstate *jump* sequence. The reader should note that the quantity coverage usually reported in the microstate literature refers to the coverage in the *continuous* sequence instead. The microstate coverage in the jump sequence can be obtained as the normalized occurrence rate of each microstate. The joint probability matrix *P*_*ij*_ is related to the conditional form of the transition matrix by *P*_*ij*_ = *T*_*ij*_ *× p*_*i*_ for *p*_*i*_ ≠0. Lehmann et al (2005) use the joint probability matrix 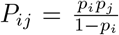 as their null hypothesis, and the conditional form is given by 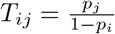.

## 3 Data and Methods

### 3.1 EEG

We analyzed n=203 eyes-closed resting-state EEG recordings from the LEMON database (Babayan et al, 2019). Preprocessed EEG files from the dataset have a sampling rate of 250 Hz and contain concatenated, non-contiguous eyes-closed and eyes-open data blocks. These were split at the indicated boundaries and epochs were analyzed further only if their duration was at least 60 seconds. In total, we obtained 1276 eyes-closed epochs from 203 subjects.

Further analyses were conducted with the Python software packages MNE and pycrostates (Gramfort, 2013; Férat et al, 2022). Eyes-closed resting-state epochs were digitally filtered with a zero-phase, non-causal band-pass filter (1-30 Hz, -6dB cutoff frequencies 0.5 Hz and 33.75 Hz), and an average reference was computed. Modified K-means clustering was performed on EEG data vectors at local peaks (temporal maxima) of the global field power time series. Modified K-means clustering was used to obtain K=4 template maps per subject in a first step, and to compute group-level microstate maps in a second clustering step (Pascual-Marqui et al, 1995; Férat et al, 2022). The group-level maps were labelled manually according to standard conventions (Michel and Koenig, 2018) and are shown in Figure A1. Continuous microstate sequences were obtained by competitive back-fitting of the group-level maps at each time point, followed by smoothing of the continuous sequence with parameters *λ* = 5 (Besag factor), and a window half-size of *b* = 3 samples, resulting in a smoothing window size of 28 ms (Pascual-Marqui et al, 1995), and a minimum segment length of 3 samples (12 ms). The first and last segment of each microstate sequence were excluded from analysis as their true duration is unknown.

### 3.2 Statistics

Markov properties of order 0, 1 and 2 were assessed with the Chi-squared tests described previously (Kullback et al, 1962; von Wegner et al, 2017). When empirical transition matrices were tested against a reference matrix with defined coefficients *T*_*ij*_, we applied the Chi-squared test defined in Kullback et al (1962). The test statistic is

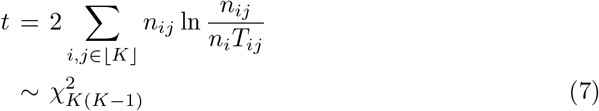

where *n*_*ij*_ are the counts of (*X*_*t*_ = *i, X*_*t*+1_ = *j*) occurrences, and *n*_*i*_ = ∑_*j*_ *n*_*ij*_ are the row sums.

#### 3.2.1 Multiple comparisons correction

When statistical tests were calculated for each microstate sequence, p-values were adjusted according to the Benjamini-Hochberg false discovery rate (FDR), at a significance level of *α* = 0.05 (Benjamini and Hochberg, 1995).

## 4 Results

### 4.1 A definition of higher-order syntax

The syntax analysis introduced in Lehmann et al (2005) analyzes the statistics of microstate words of length two. A natural extension of this concept is to consider longer microstate words 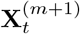, *m >* 1. The same short sequence example used above, CBACDCDBDC, can be read in blocks of words of size *m* = 4 (CBAC, BACD, etc.), for example. Higher-order syntax can then be described by either the conditional probability distribution

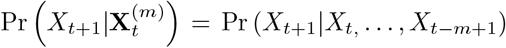

or the joint probability distribution

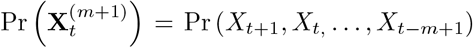

of microstate words with *m >* 1. A single element of the conditional distribution asks ‘what is the probability that the next state (*X*_*t*+1_) is C, given that the last three states were D, A, B’, for instance, and the whole distribution covers all possible label combinations. The same label combination can be described by a joint distribution, which is interpreted as a probability distribution over microstate words of length *m* = 4, in our example. There is no fundamental difference between the two descriptions since 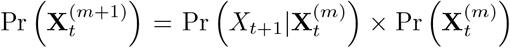, when 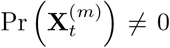, which is usually the case for microstate sequence analysis.

Starting from these distributions, many different syntax definitions can be developed. In a brute-force approach, one could enumerate all word probabilities contained in these discrete distributions but the size of these dictionaries grows exponentially with word length *m*. Instead, acknowledging the stochastic nature of these sequences, words can be grouped together, or the word distribution as a whole can be characterized by its entropy or related properties.

The general definition given above covers not only the classical syntax analysis in Lehmann et al (2005), but also studies of conditional distributions (entropy rate, active information storage, excess entropy, (von Wegner et al, 2018; Wiemers et al, 2023; von Wegner et al, 2023; Hermann et al, 2024), sample entropy as used in Murphy et al (2020), and the joint probability-based Microsynt (Artoni et al, 2023). This definition results in a hierarchy of higher-order syntax properties that only differ in the length of microstate words, or microstate histories, as shown in Table 1. We will not review these different approaches here, and instead work with the underlying conditional distribution. The question common to all approaches is how to synthesize surrogate sequences to test whether the data actually contains higher-order syntax properties (*m >* 1), independent of the syntax definition.

**Table 1.**
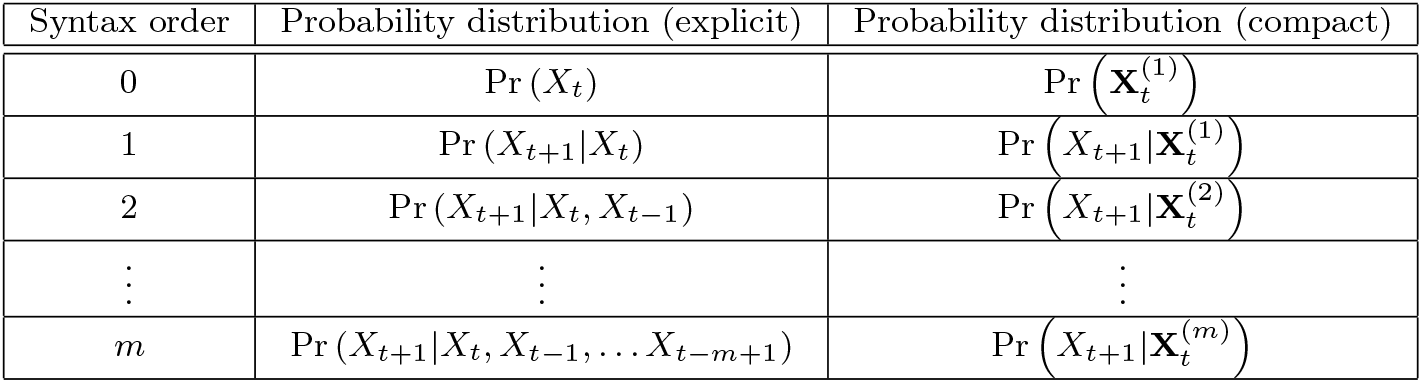
Higher-order microstate syntax, expressed as discrete conditional probability distributions. For syntax order *m* = 0, syntax is defined by the microstate distribution Pr (*X*_*t*_). For *m* = 1, syntax is defined by the conditional distribution Pr (*X*_*t*+1_ |*X*_*t*_), also known as the transition matrix of the microstate sequence. Other authors use the joint probabilities Pr (*X*_*t*_, *X*_*t*+1_) instead.

### 4.2 Requirements for first-order surrogate tests

Our aim is to obtain a statistical procedure that tests whether EEG microstate sequences have syntax properties beyond first order. The approach we follow here is to construct surrogate sequences with defined first-order properties. Computing the higher-order syntactic property of interest over these surrogates results in a null distribution against which the microstate sequences with putative higher-order properties can then be tested. To classify observed microstate word properties as higher-order syntax, i.e. higher than first order, surrogates must have the same first-order properties as the tested sequences, and must not have any properties beyond the first order, that is properties described by the transition matrix. True higher-order syntactic properties are then recognized as microstate word characteristics that cannot be predicted by extending first-order transition matrix properties to the given word length 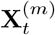,as detailed below. These considerations can be summarized in the following three requirements for surrogate microstate sequences:

**Requirement 1 (R1)** Surrogates have the same microstate distribution **p** as the original sequence.

**Requirement 2 (R2)** Surrogates have the same first-order transition matrix **T** as the original sequence.

**Requirement 3 (R3)** Surrogate data have no properties higher than first-order. This is equivalent to saying that all syntactic properties for *m >* 1 are encoded in the first-order transition matrix **T**.

Requirement **R3** implies that the probability of a given microstate word (*i*_0_*i*_1_ … *i*_*m*−1_) under first-order assumptions is given by

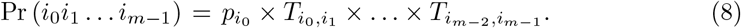

For example, the probability of the microstate word ACDBA, with time increasing from left to right, is *p*_0_ *× T*_02_ *× T*_23_ *× T*_31_ *× T*_10_, if produced by a first-order process with parameters **p** and **T**, and identifying microstate labels A-D with indices 0-3.

For convenience, we have written the microstate word with time running from left to right, while the formal microstate word expression 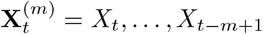 has the time indices in descending order, i.e. starting with the most recent label.

### 4.3 Generating surrogate data (1): the Markov chain method

The transition matrix *T*_*ij*_ = Pr (*X*_*t*+1_ = *j*|*X*_*t*_ = *i*) itself is a recipe for surrogate data synthesis (von Wegner and Laufs, 2018). For a microstate sequence over *K* microstate classes of length *N*, with distribution **p** and transition matrix **T**, a first-order surrogate is obtained from the following algorithm.

#### 1. Initialize

Choose the initial microstate label *s*_0_ ∈ ⌊*K*⌋ from a uniformly distributed random variable *r*_0_ ∼ 𝒰 _[0,1]_ and the microstate distribution **p** according to

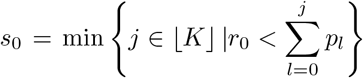

For example, assume *K* = 4, where the indices *j* = 0, …, 3 stand for microstates A, B, C, D, and a microstate distribution **p** = (0.1, 0.2, 0.4, 0.3). The initial state of the surrogate is determined by the random number *r*_0_:

a. for *r*_0_ ≤ 0.1, the initial state of the surrogate sequence is *j* = 0
b. for 0.1 *< r*_0_ ≤ 0.3, the initial state is *j* = 1
c. for 0.3 *< r*_0_ ≤ 0.7, the initial state is *j* = 2, and
d. for 0.7 *< r*_0_ ≤ 1.0, the initial state is *j* = 3

#### 2. Iterate

For *n* = 1, …, *N* − 1, generate the next microstate label *s*_*n*_ using the random number *r*_*n*_ ∼ 𝒰 _[0,1]_:

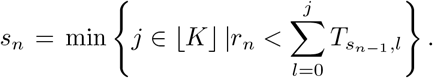

The procedure to select the next state is identical to step 1, just that the transition matrix **T** provides one probability distribution for each preceding state *s*_*n*−1_. **T** can be viewed as a family of *K* microstate label distributions *q*_*i*_ = [*T*_*i*,0_, …, *T*_*i,K*−1_], *i* ∈ ⌊*K*⌋, and *i* is selected at each time step according to the preceding state *s*_*n*−1_.

#### 4.3.1 Properties of Markov chain surrogates

Data generated with the Markov chain method follow the requirements **R1**-**R3** defined above. The theoretical fulfillment of properties **R1** and **R2** is demonstrated in (Häggström, 2002). Numerical convergence for EEG microstate sequences from the LEMON database is shown in Figure 1. The left plot shows convergence of **p**^*MC*^, the microstate distribution from Markov chain surrogates, towards the target microstate distribution **p**. The right side shows convergence of the empirical transition matrices **T**^*MC*^, obtained from Markov chain surrogates, towards **T**. Microstate distributions and transition matrices were computed at the subject level and both plots average across all coefficients *p*_*i*_ and *T*_*ij*_, respectively (mean: thick lines, 95% confidence interval: grey-shaded area).

**Fig. 1.**
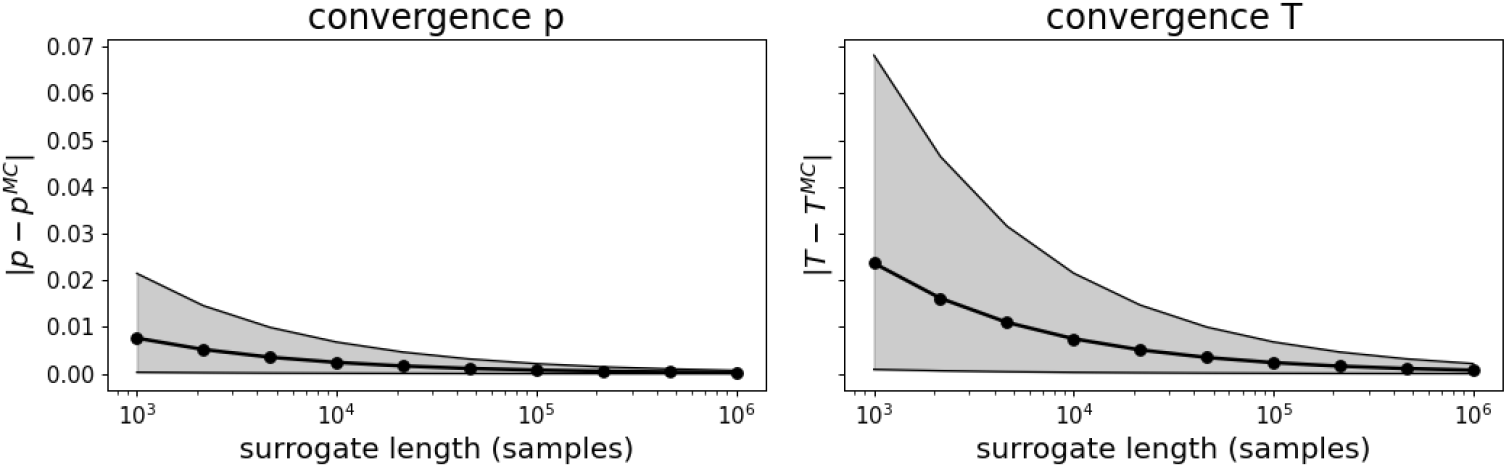
Convergence of Markov chain generated surrogate sequences towards first-order syntax. With increasing surrogate sequence length, the surrogate distribution **p**^*MC*^ and transition matrix **T**^*MC*^ asymptotically approach the nominal parameters **p** and **T**. Thick lines: mean, 95% confidence interval: grey shaded area.

Fulfillment of requirement **R3** is demonstrated by two tests. First, we applied Markov property tests of order 0, 1, and 2 to Markov chain surrogates. We found that all 1276/1276 surrogate sequences were classified as first-order processes, i.e. Markov order 0 was rejected, and orders 1 and 2 were accepted. The second test was conducted to make sure that word probabilities for word lengths *m* ≥ 3 were fully predicted by the transition matrix and compared theoretical (Equation 8) and empirical word probability distributions. The results are shown in Table 2 where the central column shows that Markov chain surrogates based on (**p, T**) produced word distributions that were indistinguishable from the expected word frequencies in most cases.

**Table 2.** Fraction of microstate word probability distributions that were significantly different from the expected first-order word probabilities (Equation 8). Surrogates were generated using either the Markov chain parameters (**p, T**) (middle column) or the shuffled jump sequence parameters (**p**^*s*^, **T**^*s*^) (right column). Statistics were FDR corrected across 100 surrogate sequences. The surrogate sequence length was 2500 samples, K=4 microstates.

| word length $m$ | $(\mathbf{p}, \mathbf{T})$ surrogates | $(\mathbf{p}^s, \mathbf{T}^s)$ surrogates |
| --- | --- | --- |
| 3 | 0.00 | 0.91 |
| 4 | 0.00 | 0.53 |
| 5 | 0.02 | 0.83 |
| 6 | 0.04 | 0.90 |

#### 4.3.2 Python and MATLAB code

The proposed Markov chain generation method is provided in Python and MATLAB, the most commonly used programming languages for microstate processing (Férat et al, 2022; Poulsen et al, 2018; Nagabhushan Kalburgi et al, 2023; von Wegner and Laufs, 2018). Both code snippets assume that the empirical distribution **p** and the transition matrix **T** in terms of conditional probabilities have been pre-computed and stored in array variables named p and T. The core functions in Python and MATLAB are given in Algorithm 1 and Algorithm 2, respectively.

##### Algorithm 1

Python code for first-order Markov surrogates. It is assumed that the microstate distribution and conditional transition matrix have been pre-computed and stored in arrays named p and T, respectively. The surrogate sequence is stored in *y* and has length *n*. NumPy imports are performed first to highlight the similarities to MATLAB code (Algorithm 2).

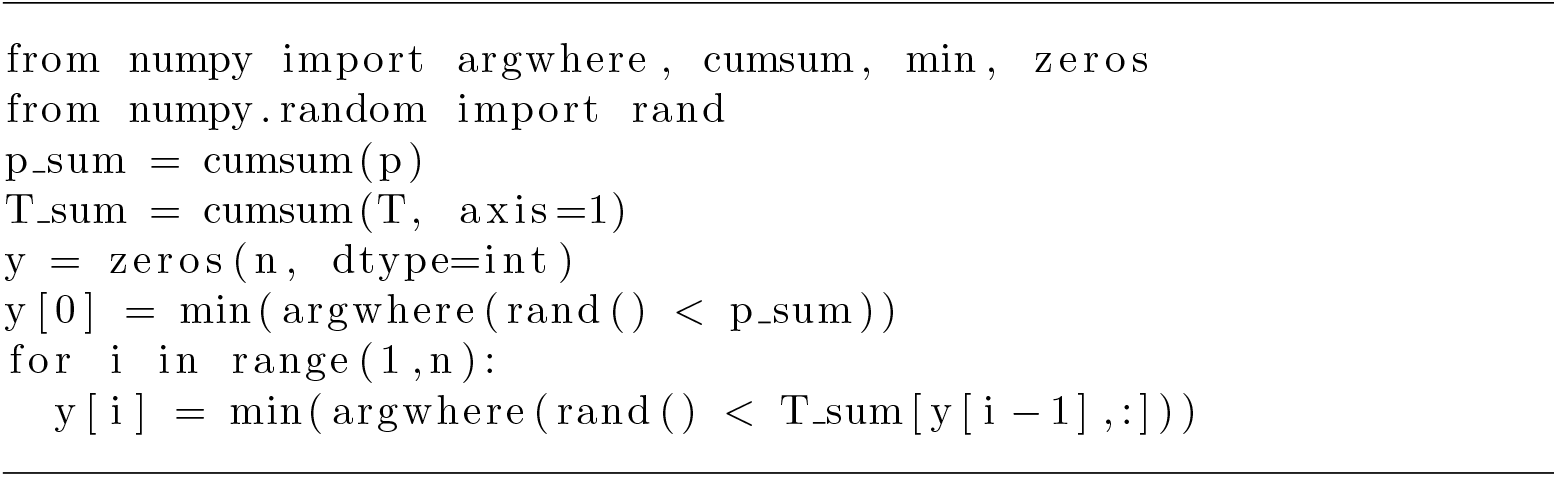

##### Algorithm 2

MATLAB code for first-order Markov surrogates. It is assumed that the microstate distribution and conditional transition matrix have been pre-computed and stored in arrays named p and T, respectively. The surrogate sequence is stored in *y* and has length *n*.

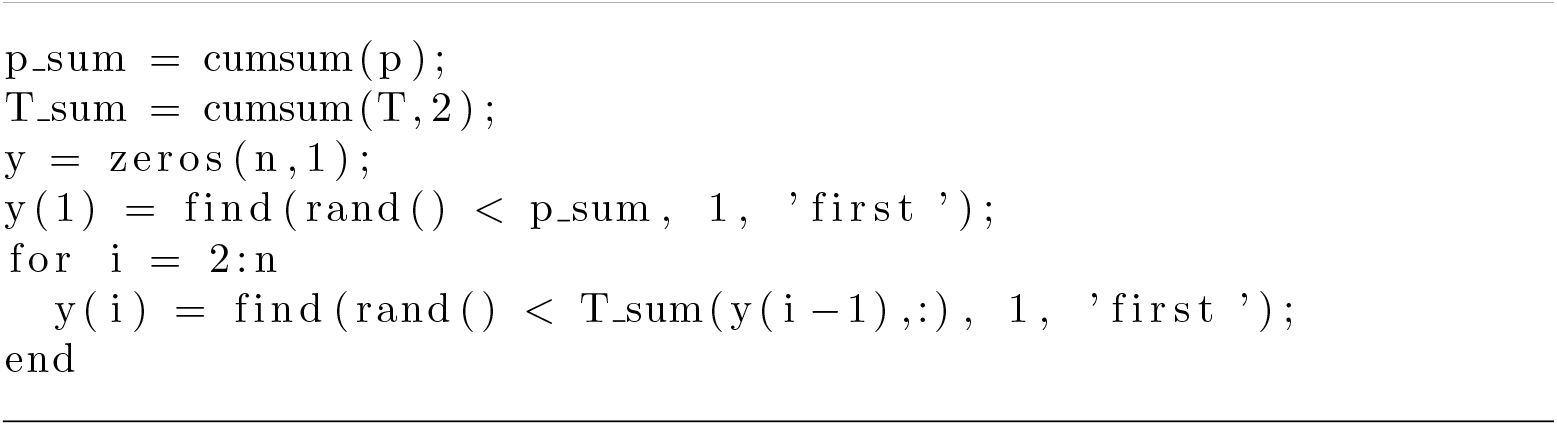

### 4.4 Generating surrogate data (2): sequence shuffling

Shuffling seeks to remove serial correlations from a time series. Computationally, it can be implemented by performing a random permutation of a given sequence, using functions such as perm in MATLAB or numpy.random.permutation in Python/NumPy. Mathematically, a perfect shuffling renders the transition probability Pr (*X*_*t*+1_|*X*_*t*_) independent of *X*_*t*_, but there exist important differences between continuous and jump microstate sequences.

#### 4.4.1 Continuous sequences

When analyzing continuous microstate sequences, i.e. sequences that may contain duplicate labels, shuffling removes all temporal correlations. Shuffled sequences are described by a zero-order transition matrix, 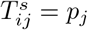 (Equation 6). In A.3, it is shown that **p** is still a valid equilibrium distribution (left eigenvector) of **T**^*s*^, i.e. shuffling does not change the microstate distribution. We tested these considerations on the LEMON dataset by generating two surrogate sequences for each continuous microstate sequence. The first surrogate was synthesized using the Markov chain algorithm with **p** and the zero-order **T**^*s*^, and the second surrogate was obtained from a random permutation of the microstate sequence. Using the zero-order and first-order Markov tests described in (von Wegner et al, 2017), we found that no real EEG microstate sequence fulfilled the zero- or first-order Markov property, indicating higher-order properties. For both surrogate types, Markov chain surrogates and shuffled sequences, all samples were classified as zero-order (uncorrelated) sequences. These analyses included multiple-comparisons correction (FDR).

#### 4.4.2 Jump sequences

Next, we analyzed jump sequences which are commonly used for syntax analyses. Surrogate techniques need to take into account that the resulting surrogate must not have two identical consecutive microstate labels. In the microstate literature, two different methods can be found to achieve this. The first approach, used in Artoni et al (2023), performs shuffling on the continuous microstate sequence, by moving contiguous blocks of microstate labels before removing duplicate labels. The second approach computes the jump sequence from the continuous sequence first, and then shuffles the labels on the jump sequence, avoiding duplicates in the process. Figure 2 illustrates both approaches in a commutative diagram that shows the equivalence of both approaches. The following analysis is therefore valid for both approaches.

**Fig. 2.**
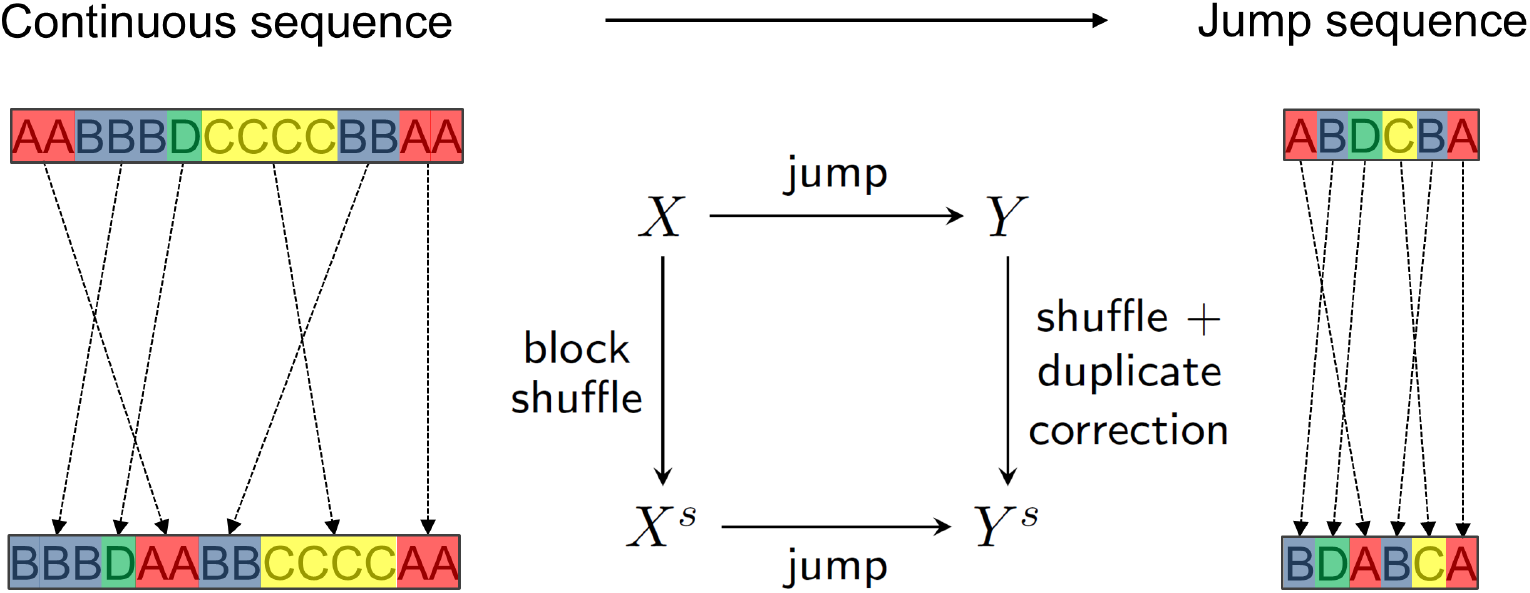
Two equivalent shuffling approaches for microstate jump sequences. In this diagram, *X* denotes a continuous EEG microstate sequence, and *Y* the embedded jump sequence. *X*^*s*^ and *Y* ^*s*^ denote the shuffled continuous and jump sequence, respectively. First approach: Shuffling of contiguous blocks of microstate labels in the continuous microstate sequence (*X* →*X*^*s*^, block shuffle) precedes the computation of the jump sequence (*X*^*s*^ →*Y* ^*s*^). Second approach: computing the embedded jump sequence (*X* →*Y*) is followed by shuffling, while preserving the jump property by removing duplicates (*Y*→ *Y* ^*s*^, shuffle + duplicate-correction). The diagram in the center of the image is commutative, showing the equivalence of both approaches. The first approach has been used in (Artoni et al, 2023), and the second approach leads to the transition matrix introduced by Lehmann et al (2005). A slight variation of the second method has been used in Murphy et al (2020).

#### 4.4.3 Jump sequence shuffling changes the equilibrium distribution

Shuffling the microstate jump sequence with transition matrix **T** and distribution **p** yields a preliminary transition matrix **T**^*l*^ with coefficients 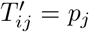, effectively decorrelating the sequence (Equation 6). The microstate distribution **p** of the original jump sequence is still a valid equilibrium distribution of **T**^′^, as shown in Appendix-A.4. However, since **T**^′^ does not describe a microstate jump sequence (diagonal elements are not zero), the transition matrix needs to be transformed by (i) setting diagonal entries to zero, and (ii) re-scaling all matrix elements to give row sums of one. These transformations are the mathematical equivalent of the processes illustrated in Figure 2. The final transition matrix of the shuffled and duplicate-corrected jump sequence is

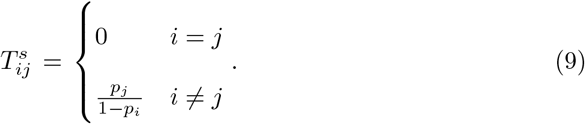

This is the conditional form of the null hypothesis derived in Lehmann et al (2005), demonstrating that the approaches used in Lehmann et al (2005) and Artoni et al (2023) are equivalent. However, the microstate distribution **p** of the original jump sequence is not an eigenvector of **T**^*s*^, as shown in A.4. Thus, the shuffling procedure described in Figure 2 produces sequences whose microstate distribution is not the same as the sequence under investigation. This property contradicts our surrogate requirement **R1**. The left side of Figure 3 shows the distribution of the absolute deviations 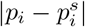 between the original microstate probabilities (*p*_*i*_) and their shuffled/duplicate-corrected surrogates 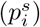. The distribution is pooled over all indices *i* ∈ ⌊*K*⌋, and all n=1276 microstate jump sequences from the LEMON database. The mean deviation was 0.01. The distribution of the original jump sequences was **p** = (0.21, 0.22, 0.27, 0.30).

**Fig. 3.**
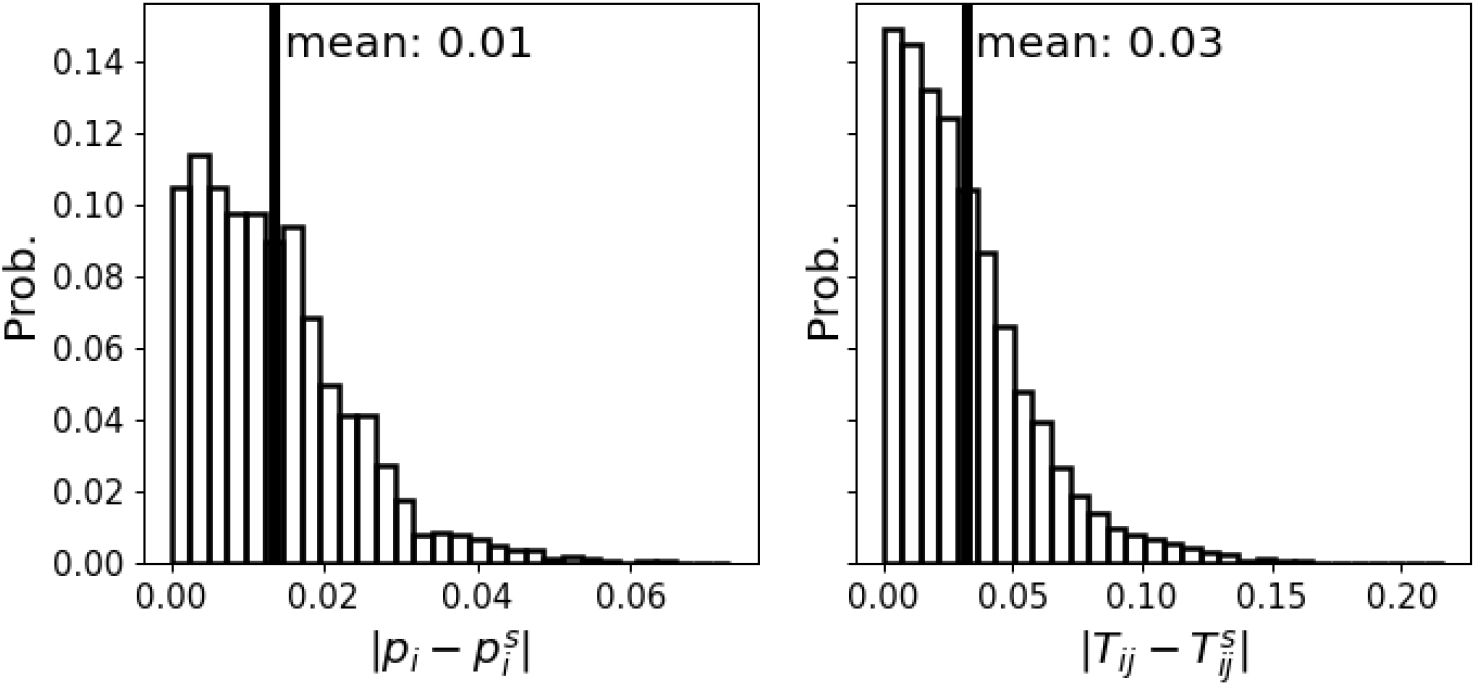
Distribution of absolute parameter deviations between original microstate jump sequences and shuffled surrogate sequences. Systematic deviations are shown for microstate distributions 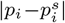 (left), and for transition matrix coefficients 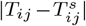 (right), pooled over all *i, j* ∈ ⌊*K* ⌋ and all n=203 subjects in the LEMON database. Vertical solid lines indicate the mean of the distribution.

#### 4.4.4 Jump sequence shuffling changes the transition matrix

This section analyses the differences between transition matrix coefficients from original microstate jump sequences (*T*_*ij*_) and shuffled/duplicate-corrected surrogates 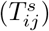. The right side of Figure 3 shows the distribution of the deviations 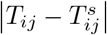, pooled across all indices *i, j* ∈ ⌊*K*⌋ and all microstate jump sequences. The mean deviation was 0.03 and 45% of all deviations were larger than 0.03.

A concrete example from a single subject is shown in Figure 4. The left side shows the actual transition matrix **T** of a microstate jump sequence, and the matrix in the center is the corresponding transition matrix produced by shuffling/duplicate-correction (**T**^*s*^). On the right, the difference **T** − **T**^*s*^ is shown. The largest deviations in either direction are -0.12 and 0.22. These findings contradict our surrogate requirement **R2**.

**Fig. 4.**
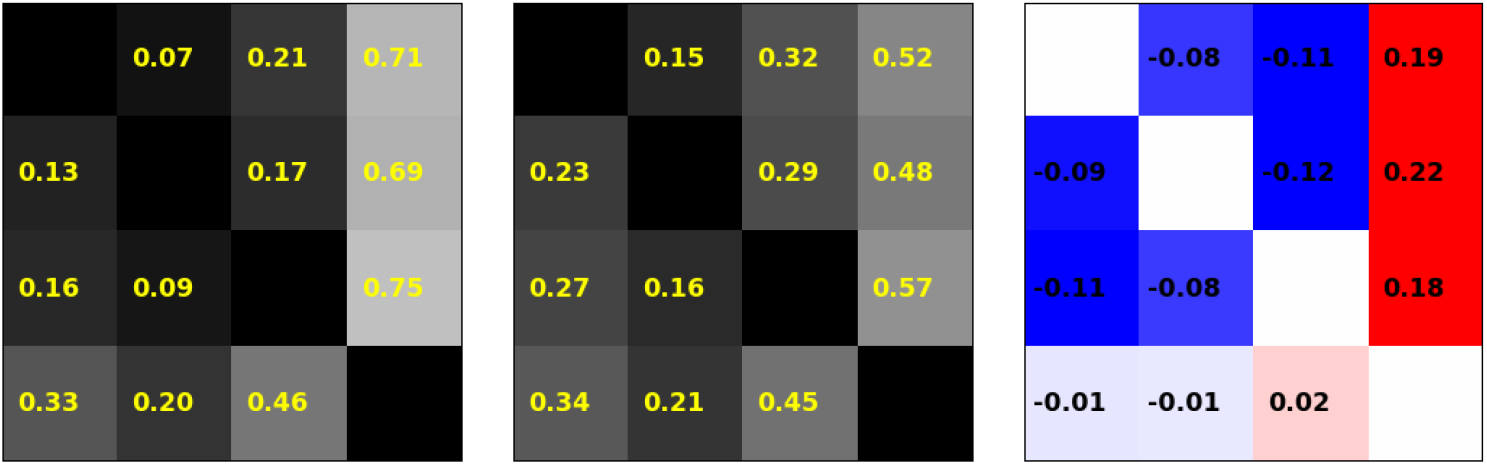
The effects of shuffling on microstate transition matrices. **A**: Transition matrix **T** of the original microstate jump sequence in an eyes-closed resting state. **B**: Transition matrix **T**^*s*^ of the shuffled jump sequence. **C**: Difference between **A** and **B**.

#### 4.4.5 Shuffling produces false positive higher-order syntax results

The previous findings show that jump sequence shuffling changes both the microstate distribution **p** and the transition matrix **T**. Next, we wanted to understand to which extent the differences between the true first order Markov chain parameters **p** and **T** and the shuffled sequence parameters (**p**^*s*^, **T**^*s*^) would affect microstate word probabilities. We know that surrogates generated from (**p, T**) should converge towards these expected values and that those generated by (**p**^*s*^, **T**^*s*^) will eventually be different, but it is not clear whether these differences will be noticeable in empirical microstate word distributions.

We therefore compared expected and observed word probabilities in microstate jump sequences generated from each parameter set. First, we calculated a single microstate probability distribution **p** and transition matrix **T** across all n=203 subjects from the LEMON dataset. From both parameter sets, (**p, T**) and (**p**^*s*^, **T**^*s*^), we generated 100 surrogate sequences with a length of 2500 samples each and compared the resulting microstate word probabilities with those expected from first-order properties (**p, T**) alone. The expected word probabilities under parameters (**p, T**) are given by Equation 8.

We tested word lengths *m* = 3, …, 6, for which there are *n*_*m*_ = *K ×*(*K* − 1)^*m*−1^ words without microstate duplicates. We compared expected and observed microstate word distributions with a Chi-squared test and performed FDR multiple comparisons correction over the 100 p-values obtained from the 100 surrogate sequences. The results are shown in Table 2, where the fraction of significantly different word probability distributions is indicated for Markov chain surrogates (parameters (**p, T**), middle), and for shuffled surrogates (parameters (**p**^*s*^, **T**^*s*^), right). It is observed that almost all word distributions from shuffled sequences (**p**^*s*^, **T**^*s*^) were measurably different from the distributions expected under (**p, T**).

#### 4.4.6 Jump sequence shuffling leads to shorter surrogates

Shuffling can cause identical microstate labels to end up next to each other. The reformatting into a correct jump sequence requires deletion of these duplicate labels and shortens the output sequence. For example, if ABDCBA is shuffled into BBCAAD, the duplicates have to be removed and the output sequence BCAD is two samples shorter than the input sequence. This effect does not occur with the Markov chain generation method as the diagonal of the transition matrix has zeros and the probability of generating a duplicate pair (CC, or similar) is zero. The extent of this effect depends on the average microstate duration and occurrence. In the LEMON dataset, we found microstate durations of 36.9 ms (microstate class A), 37.4 ms (B), 40.1 ms (C), 45.4 ms (D), and occurrence values of 5.2/s (A), 5.3/s (B), 6.7/s (C), and 7.3/s (D). The observed effect was an average sequence shortening from 1678 (1354,2363) to 1240 (964,1754) samples (mean, 95% confidence intervals). On average, the ratio of sequence lengths after to before shuffling was 0.74 (0.70,0.76), so approximately one quarter of the data was lost.

#### 4.4.7 The Murphy et al. 2020 algorithm

The jump sequence shuffling method published in Murphy et al (2020), along with its MATLAB implementation, is not a pure sequence shuffling algorithm but contains extra steps to avoid duplicates. Since these additional steps cannot be modelled as a transition matrix, we used the published method to compute surrogates for the LEMON database microstate jump sequences and tested their transition matrices against the original transition matrix **T**, and against the theoretical matrix for shuffled sequences **T**^*s*^ described in Equation 9, using the test described in Equation 7. After FDR correction, 64% of all surrogates had transition matrices that were significantly different from **T**, and 74% were significantly different from **T**^*s*^.

## 5 Discussion

This study was motivated by a growing interest in multivariate EEG microstate statistics and how these might generate new insights into brain function and pathology (Michel and Koenig, 2018). The notion of microstate syntax has been used for years (Lehmann et al, 2005), focusing on microstate words of length two, i.e. bivariate distributions, expressed as either conditional or joint probabilities. Given the larger number of studies using this approach, our understanding of bivariate distributions is more advanced than that of higher-order multivariate distributions (Michel and Koenig, 2018).

The principal aim of this study was to investigate two methods for surrogate data generation regarding their usefulness to detect higher-order syntax in microstate sequences. While our aim was not to introduce or evaluate a specific higher-order syntax metric, we found it necessary to provide a definition of higher-order syntax in order to define the scope of the surrogate data methods discussed here.

We propose to use a natural extension of the existing first-order syntax definition to higher orders by using multivariate conditional or joint probability distributions, being aware that there is no fundamental difference between the conditional and joint forms. In our interpretation, any method that quantifies either 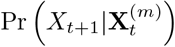 or 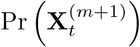 for *m >* 1 represents a higher-order syntax analysis. An advantage of this broad definition is that many different syntax definitions can be derived from these distributions, and thus, the surrogate statistical methods discussed here are expected to be useful not only for currently used syntax concepts, but also for future developments. In recent years, we have used Shannon entropy to characterize conditional distributions for different word lengths (von Wegner et al, 2017, 2018, 2023; Wiemers et al, 2023; Hermann et al, 2024; Jia et al, 2021), using metrics known as entropy rate, excess entropy, and active information storage (Lizier et al, 2012). Other authors have chosen other entropy variants such as sample entropy (Murphy et al, 2020), or have pooled microstate words into classes of similar word entropy (Artoni et al, 2023). All the aforementioned techniques have chosen approaches that compress the information contained in the full multivariate distributions. We believe that this approach is preferable to an exhaustive (or ‘brute-force’) approach that enumerates the whole microstate word dictionary, i.e. all possible microstate words and their probabilities. Microstate word dictionary size grows exponentially with word length *m*, which makes the practical interpretation of full dictionaries difficult. While others syntax definitions can be conceived, e.g. methods from the theory of formal languages and automata (Grassberger, 1988), we believe that methods based on word probability distributions will play an important role in the near future. Any such method would fall under the definition used in this article.

What has been achieved in this study is an in-depth analysis of surrogate sequences obtained from two different methods, namely Markov chain generated surrogates and surrogates from sequence shuffling, both of which have been used in the microstate literature (von Wegner et al, 2016, 2017; Wiemers et al, 2023; Hermann et al, 2024; von Wegner et al, 2023; Murphy et al, 2020; Artoni et al, 2022, 2023).

The following is a summary of our results:

1. An adequate null hypothesis to test higher-order microstate syntax is the sequence’s own first-order syntax, described by the microstate distribution and transition matrix.
2. This leads to three requirements for surrogate sequences: they should have the same microstate distribution **p** and the same first-order transition matrix **T** as the original sequence (**R1, R2**), and no temporal dependencies higher than first order (**R3**), i.e. all higher-order word properties of these surrogates must be predictable from **p** and **T**.
3. Microstate jump sequences have at least first-order structure. The jump condition (no duplicates) is incompatible with an uncorrelated, zero-order surrogate structure. For continuous microstate sequences, however, zero-order surrogates can be obtained via the Markov chain or the shuffling method.
4. Surrogate requirements **R1**-**R3** are fulfilled for Markov chain surrogates, i.e. when surrogates are generated from the sequence’s transition matrix.
5. Surrogate requirements **R1**-**R3** are not fulfilled when jump sequences are randomly shuffled and then corrected for duplicate labels. The latter leads to the same transition matrix that provides the null hypothesis in Lehmann et al (2005).
6. Shuffling/duplicate-correction produces surrogates that are up to 25% shorter than the original sequence in our dataset, reducing the power of statistical tests.

### 5.1 Markov chain generation

We have given a simple computational implementation of the Markov chain generation method that automatically fulfills surrogate requirements **R1**-**R3**. The algorithms have a predictable running time that depends linearly on the desired length of the output sequence (𝒪 (*n*)), as opposed to other microstate shuffling algorithms (Murphy et al, 2020). The Markov chain method can be used to generate surrogates of arbitrary length. This is not possible when using shuffling/duplicate-correction which produces surrogates that are even shorter than the input sequence, or exact shuffling approaches that always reproduce the exact input length (Murphy et al, 2020). Generating surrogates with more samples than the input sequence can be of interest when investigating syntax measures for which theoretical values are not available. Entropy rate values, for example, can be computed analytically from a Markov chain’s distribution and transition matrix (von Wegner et al, 2018), but no theoretical distribution of entropy classes as used in Microsynt (Artoni et al, 2023) is available at the moment. In these cases, the correct asymptotic distribution of the syntax metric under first-order assumptions can be approximated with surrogates, and moreover, the numerical behaviour of the metric as a function of sample size can be measured. We have used this approach to study how Lempel-Ziv complexity and entropy rate values converge as a function of microstate sequence length, using far greater sequence lengths than available from experiments (von Wegner et al, 2023).

### 5.2 Shuffling

Sequence shuffling is an intuitive approach to remove temporal correlations from a time series. When it comes to microstate sequences, and jump sequences in particular, shuffling and post-shuffling procedures imprint certain properties on the generated surrogate sequences.

The simplest case is that of a continuous microstate sequence, i.e. the sequence obtained from competitively back-fitting the microstate templates to each time point of the EEG dataset, possibly followed by certain post-processing steps such as smoothing. The resulting continuous sequence will have many consecutive duplicate labels, given the relatively long lifetime of a microstate (tens to hundreds of milliseconds) compared to the EEG sampling rate (2-10 ms). For these sequences, computing a simple permutation of the sequence gives a zero-order Markov surrogate. This approach has been used, among other tests, to search for long-range correlations in microstate sequences (Van de Ville et al, 2010). Zero-order surrogates can be used to test for the presence of a first-order syntax. If the null hypothesis is not rejected, the tested sequence would be classified as completely random. We are not aware of any experimental condition for which this has been reported for EEG microstate sequences.

An important observation is that if a zero-order surrogate of a continuous sequence is transformed into a jump sequence, by removal of all consecutive duplicate labels, the resulting jump sequence is lifted to a first-order syntax. The reason for this is that being a jump sequence and being uncorrelated are incompatible properties. An uncorrelated sequence with microstate distribution **p** = (*p*_*i*_)_*i*∈⌊*K*⌋_ has transition matrix structure *T*_*ij*_ = *p*_*j*_ (every row is **p**), but being a jump sequence requires *T*_*ii*_ = 0 for all *i* ∈ ⌊*K*⌋ (no self-transitions). This is impossible as it implies *p*_*i*_ = 0 for all *i* ∈ ⌊*K*⌋. Thus, the decorrelating property of the shuffling procedure is partially reversed by the removal of duplicate labels. The jump sequence obtained from a random continuous sequence has a syntax order (*m* = 1) that is higher than the continuous sequence from which it has been obtained (*m* = 0). Moreover, an inverse inference is not possible. If a jump sequence has exactly the syntax that is expected from a zero-order continuous sequence, it cannot be inferred that the continuous sequence actually was completely random. Counterexamples are easily constructed: starting with the jump sequence, and interpolating any microstate lifetime distribution in-between the microstate switches, one would obviously obtain the same jump sequence back when removing the duplicates. The continuous sequence from which the jump sequence was obtained could therefore have short-range correlated Markov features, long-range correlated (fractal) properties, or any other correlation structure. Information about the correlation structure of the continuous sequence is lost when transitioning to the jump sequence. Although shuffling might be suggestive of removing all correlations, a sequence will at least have first-order properties as long as the jump sequence property is fulfilled.

Our results show that the requirements **R1**-**R2** are not fulfilled by shuffled/duplicate-corrected surrogates. The requirements are met in the exceptional case of a uniform microstate distribution, but not in general. Interestingly, shuffling a continuous microstate sequence leads to a transition matrix 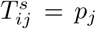 that is not changed when the shuffling step is applied again, since **p** is an equilibrium distribution of **T**^*s*^. The same is not true for jump sequences, where repeated application of shuffling/duplicate-correction converges towards an entirely different equilibrium distribution and transition matrix, namely 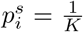, and 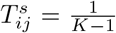 for the off-diagonal elements of the transition matrix. There is no obvious reason to test an arbitrary microstate sequence against such a null hypothesis that is fundamentally different from the observed microstate sequence, but the shuffling/duplicate-correction method moves the surrogates one step into the direction of this oversimplified model. Nevertheless, all models created along that path will have first-order syntax. We have not found an explicit discussion of these relationships in the existing microstate literature.

#### 5.2.1 Microstate distributions and transition matrices

For jump sequence surrogates obtained from shuffling/duplicate-correction, we observed changes in the microstate distribution **p** (coverage) and the transition matrix **T**, when compared to the underlying sequence. The mean absolute deviation of coverage values 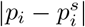 was 0.01, which might appear to be a small error, relative to the absolute coverage values of 0.19-0.34. Nevertheless, the resulting microstate word probabilities were significantly different, as discussed below. Artoni et al (2023) discussed an alteration of the microstate distribution after shuffling and chose a larger number of surrogates to account for the effect. Our analysis however shows that this is a systematic deviation. A larger number of surrogates will generate the altered distribution **p**^*s*^ with higher precision, but will not correct the systematic deviation from **p** (Figure 3), which is fully encoded in **p** itself (A.4).

For transition matrix entries, the mean absolute deviation 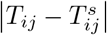 was 0.03. Almost half (45%) of the deviations were larger than 0.03, and reached a maximum value of 0.22. The mean deviation already lies within the range of effect sizes reported in microstate syntax studies, and the larger deviations clearly exceed these. Al Zoubi et al (2019) used conditional probabilities for first-order syntax analysis of jump sequences and found significant changes in individual transition rates ranging between 0.002 and 0.045. Tomescu et al (2015) analyzed continuous sequences and found significant conditional transition probability differences into microstate class C in schizophrenia patients, with differences of approximately 0.02-0.03. They also reported increased B-to-C transition probabilities in 22q11 deletion syndrome individuals, with a difference in mean probabilities of 0.024. A later study by the same group used Lehmann et al (2005) syntax analysis, and some of the significant transition probability differences were below 0.05 (Tomescu et al, 2018). An et al (2024) used joint probabilities of jump sequences and found rate differences mostly with the range -0.03, 0.03. Nishida et al (2013) used joint transition probabilities of jump sequences and the maximal difference across all group combinations was 0.02.

#### 5.2.2 Word probabilities

While the differences in distribution and transition matrices were sufficient to show that shuffled surrogates do not fulfill **R1**-**R2**, we were interested in measuring the effect of these deviations in **p** and **T** on actual microstate word distributions, as these could have been negligible after all. We found that the difference between first-order properties of shuffled sequences and the original sequence are more than just a numerical inaccuracy in analytical expressions and eigenvector equations. Almost all word probability distributions obtained from shuffled surrogates were statistically different from the original sequences’ first-order properties, contradicting our surrogate requirement **R3** (Table 2). It is important to highlight that this does not reflect a stronger decorrelation of shuffled sequences. Both, shuffled and Markov chain-generated sequences have a first-order Markov structure, but their parameters (distribution **p**, transition matrix **T**) are different.

This could easily lead to false positive results in higher-order syntax analyses. Assume that a given microstate jump sequence is a perfect first-order Markov chain, without higher-order properties. To test this sequence for higher-order syntax, we could compare the microstate word probabilities contained in the sequence to a set of shuffled jump sequences, and we would find statistically significant differences with very high probability (see Table 2). Since we know that the shuffled sequences have first-order structure, and there is a difference between the test sequence and the surrogates, the false conclusion could be that the test sequence has higher-order syntax properties. The true reason, however, is that the difference in word probability was caused by differences between (**p, T**) and (**p**^*s*^, **T**^*s*^).

### 5.3 The Murphy et al. (2020) method

The MATLAB algorithm published in Murphy et al (2020) is described as a “random permutation of the microstate sequence labels such that the same labels could not be adjacent”. It should be noted, however, that the implementation of the algorithm contains more steps than a simple permutation procedure. The first K-1 microstates are assigned random locations in the surrogate array, while avoiding adjacent duplicates. The last microstate label is then placed into the remaining empty array elements. In most cases, this step produces adjacent duplicate labels. The algorithm then tries to remove these by performing swaps with neighbouring labels. An iterative search procedure scans for swap options, starting with labels three samples into the future of the duplicate, and widening the search up to a maximum of 20 samples if no swap candidates are found nearby. This search procedure deviates from a random permutation approach as it introduces fixed parameters and a systematically widening but limited search window for swap candidates, possibly adding short-range dependencies. Sometimes, the approach is not successful, and the current permutation attempt is abandoned. When successful, a correct jump sequence is produced, but in most cases its transition matrix is different from both, the original transition matrix of the jump sequence **T**, and from the matrix **T**^*s*^ expected from pure shuffling/duplicate-correction (Equation 9). These surrogates therefore do not comply with our requirement **R2**. As some permutation attempts fail, the algorithm has variable and unpredictable computation times. For almost 50% of the sequences in our dataset, the published algorithm could not find a valid permutation at all.

An advantage of the algorithm is that it does not change the length of the sequence, as it happens with block-shuffling (Artoni et al, 2022). Another feature of the algorithm is that each surrogate contains exactly the same copy number of each microstate as the original sequence. This feature can be seen as a strength, but also as a shortcoming of the method. If microstate sequences are regarded as sample paths generated by a stochastic process, we would not expect identical microstate copy numbers in each sample trajectory. In that sense, the Markov chain method reproduces the stochastic properties of the assumed underlying process more correctly. The expected coverage is produced exactly in the limit of many sample paths, but not reproduced exactly in individual sample paths.

### 5.4 Lehmann’s syntax analysis

The syntax analysis presented in Lehmann et al (2005) is based on the same jump sequence transition matrix (**T**^*s*^) that is obtained by block shuffling/duplicate-correction. The comparison between observed and expected joint probabilities as described in Lehmann et al (2005) and implemented in some microstate toolboxes (Nagabhushan Kalburgi et al, 2023) is therefore fundamentally different from the premises formulated in Requirements **R1** and **R2**. Statistically significant differences obtained from this test cannot be interpreted as higher-order syntactic properties. Moreover, the description “if transitions from a preceding state into a next state occurred randomly, i.e. independent of the class of the preceding state” (Lehmann et al, 2005) is easily misunderstood. Temporal independence could occur in continuous microstate sequences, but not in jump sequences, but the latter are the objects of interest in Lehmann et al (2005). Independence is reflected by a transition matrix with identical rows, however, this is not possible for a jump process, as explained above. Independence of the preceding state is lost through the correction term 1 − *p*_*i*_ in 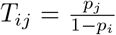 (or 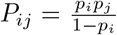 in Lehmann et al (2005)), because the index *i* refers to the preceding microstate, *T*_*ij*_ = Pr (*X*_*t*+1_ = *j*|*X*_*t*_ = *i*).

We therefore suggest to use direct tests of Markovianity to assess whether a given microstate sequence has low-order Markov properties (von Wegner et al, 2017). Jump sequences do not need to be tested for the zero-order Markov property, as this is impossible for a jump sequence. First-order Markovianity tests should be performed because a microstate sequence could be a first-order Markov process, regardless of Lehmann’s syntax test results.

### 5.5 Conclusion

We conclude that the concept of higher-order microstate syntax encompasses many different approaches, and we have given a definition that covers definitions based on microstate word distributions, either in their conditional or joint probability form. We have formulated three requirements that surrogate data should fulfill to serve as a null distribution for higher-order syntax statistical tests. A test based on such a null distribution would test the null hypothesis of first-order syntax. Quantitative analysis shows that Markov chain generation is an efficient (𝒪 (*N*)) and effective method that fulfills all requirements, whereas different forms of sequence shuffling can fail these requirements.

## 6 Conflict of interest

The authors declare that they have no conflict of interest.

**Fig. A1.**
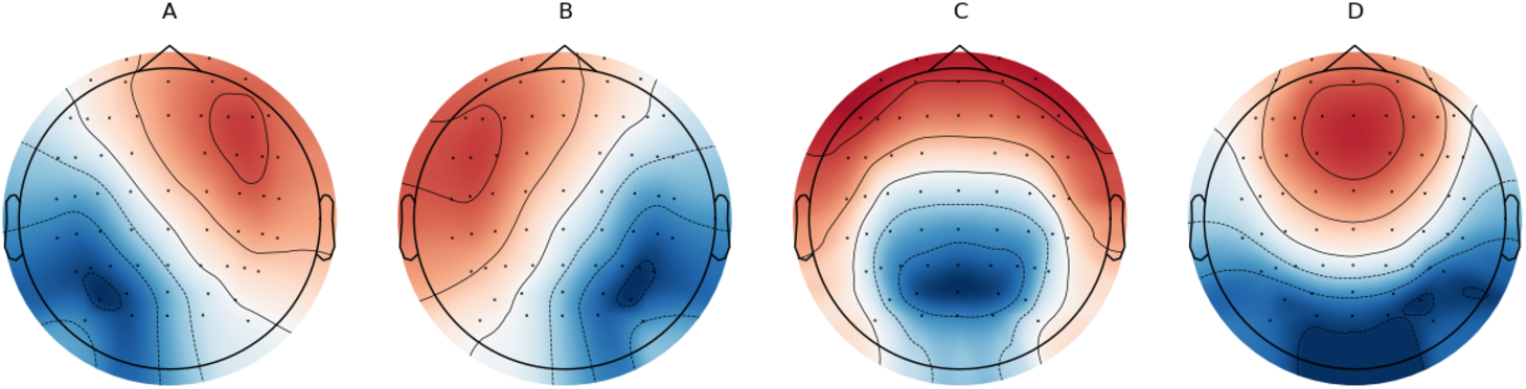
Group-level microstate maps from the modified K-means algorithm. Subject-level maps were obtained from n=203 eyes-closed resting-state EEG recordings from the LEMON dataset.

## A Appendix

### A.1 Microstate analysis

This section presents the basic results of the microstate analysis of the LEMON dataset. Microstate maps for K=4 are shown in Figure A1 and the elementary microstate statistics duration, occurrence, and coverage are given in Table A3. It should be noted that the qualitative results in the main text are not dependent on the number of clusters *K*, or the exact numerical outcomes of the microstate clustering and fitting algorithm.

**Table A3.**
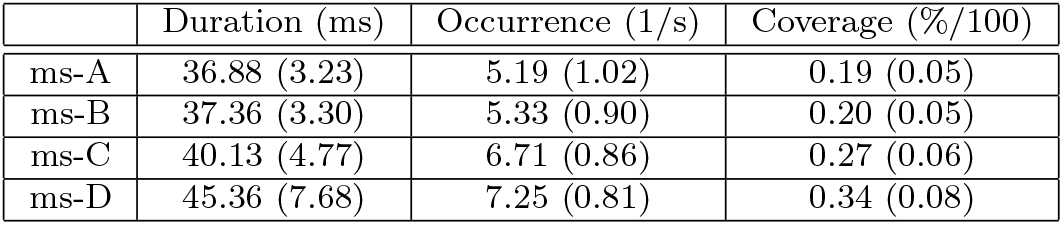
Microstate statistics as means (standard deviation) across n=203 eyes-closed resting-state EEG recordings from the LEMON database.

#### A.2 Empirical distribution and transition matrix

Using the maximum likelihood estimators for the microstate distribution **p** and the transition matrix **T** (Anderson and Goodman, 1957), we find

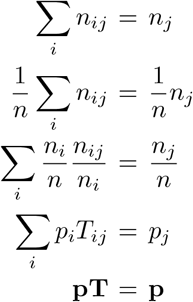

where *n*_*ij*_ is the number of co-occurrences (2-words) (*X*_*t*_ = *i, X*_*t*+1_ = *j*), *n*_*j*_ is the number of *X*_*t*_ = *j* counts, and *n* is the length of the sequence. Therefore, the empirical distribution **p** is always a left eigenvector of **T** to the eigenvalue *λ*_0_ = 1, i.e. an equilibrium distribution.

### A.3 Shuffling the continuous sequence preserves the equilibrium distribution

Consider a continuous microstate sequence with distribution **p** = (*p*_0_, …, *p*_*K*−1_) and transition matrix **T**. By construction, we have **pT** = **p** (A.2). Sequences obtained from shuffling the continuous sequence are described by the transition matrix for the corresponding zero-order process

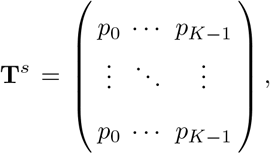

or simply 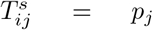, which explains the absence of correlations 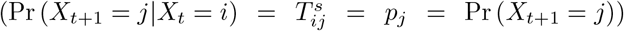. **p** is still an equilibrium distribution of the shuffling matrix **T**^*s*^. Let **q** =**pT**^*s*^, then

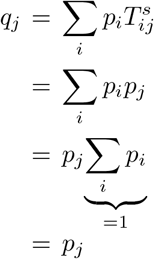

so **p** =**pT**^*s*^.

### A.4 Shuffling the jump sequence changes the equilibrium distribution

Consider a jump microstate sequence with distribution **p** = (*p*_0_, …, *p*_*K*−1_) and transition matrix **T**. By construction, we have **pT** = **p** (A.2). Sequences obtained from shuffling the jump sequence have a transition matrix **T**^′^ with coefficients 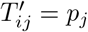:

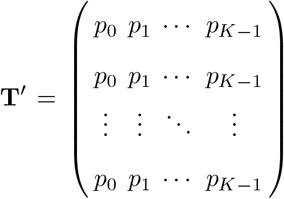

The distribution **p** is still a valid equilibrium distribution for **T**^′^, following the same arguments used in A.3. However, **T**^′^ does not describe a jump sequence as the diagonal entries are not zero. To correct for this, and to recover an admissible jump sequence matrix, two transformation must be applied, (i) the diagonal of **T**^′^ must be set to zero to disallow microstate repeats

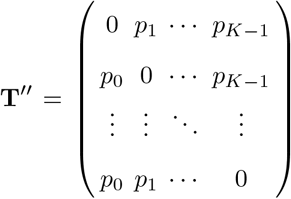

and (ii) the rows of **T**^*ll*^ must be re-normalized to give a matrix **T**^*s*^ such that 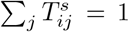, in order to obtain a stochastic matrix. The normalization constant is obtained from 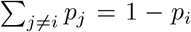, and the final transition matrix of the shuffled and duplicate-corrected sequences is

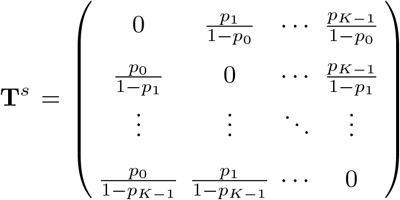

or simply

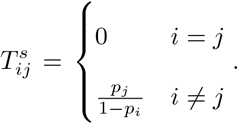

To check whether **p** is also a stationary distribution of **T**^*s*^, let **q** = **pT**^*s*^. We get

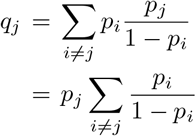

which fulfills *q*_*j*_ = *p*_*j*_ only when 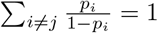. This is fulfilled by 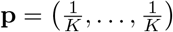 and 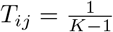 for *i* ≠ *j*, a fixed point if the shuffling/duplicate-correction procedure is iterated, but not for general **p** and **T**. Thus, shuffling of jump sequences produces first-order sequences whose first-order properties are different from the starting parameters **p** and **T**, in contradiction to requirements **R1** and **R2**.

